# The Sequence Basis for Selectivity of Ephrin-B2 Ligand for Eph Receptors and Pathogenic Henipavirus G Glycoproteins

**DOI:** 10.1101/2023.04.26.538420

**Authors:** Krishna K. Narayanan, Moushimi Amaya, Natalie Tsang, Randy Yin, Alka Jays, Christopher C. Broder, Diwakar Shukla, Erik Procko

## Abstract

Ephrin-B2 (EFNB2) is a ligand for six Eph receptors in humans and functions as a cell entry receptor for several henipaviruses including Nipah virus (NiV), a pathogenic zoonotic virus with pandemic potential. To understand the sequence basis of promiscuity for EFNB2 binding to the attachment glycoprotein of NiV (NiV-G) and Eph receptors, we performed deep mutagenesis on EFNB2 to identify mutations that enhance binding to NiV-G over EphB2, one of the highest affinity Eph receptors. The mutations highlight how different EFNB2 conformations are selected by NiV-G versus EphB2. Specificity mutations are enriched at the base of the G-H binding loop of EFNB2, especially surrounding a phenylalanine hinge upon which the G-H loop pivots, and at a phenylalanine hook that rotates away from the EFNB2 core to engage Eph receptors. One EFNB2 mutant, D62Q, possesses pan-specificity to the attachment glycoproteins of closely related henipaviruses and has markedly diminished binding to the six Eph receptors. However, EFNB2-D62Q has high residual binding to EphB3 and EphB4. A second deep mutational scan of EFNB2 identified combinatorial mutations to further enhance specificity to NiV-G. A triple mutant of soluble EFNB2, D62Q-Q130L-V167L, has minimal binding to Eph receptors but maintains binding, albeit reduced, to NiV-G. Soluble EFNB2 decoy receptors carrying the specificity mutations were potent neutralizers of chimeric henipaviruses. These findings demonstrate how specific residue changes at the shared binding interface of a promiscuous ligand (EFNB2) can influence selectivity for multiple receptors, and may also offer insight towards the development of henipavirus therapeutics and diagnostics.

**IMPORTANCE:** Ephrin-B2 (EFNB2) is a ligand for six Eph receptors in humans and regulates multiple cell developmental and signaling processes. It also functions as the cell entry receptor for Nipah virus and Hendra virus, zoonotic viruses that can cause respiratory and/or neurological symptoms in humans with high mortality. Here, we investigate the sequence basis of EFNB2 specificity for binding the Nipah virus attachment G glycoprotein over Eph receptors. We then use this information to engineer EFNB2 as a soluble decoy receptor that specifically binds the attachment glycoproteins of Nipah virus and other related henipaviruses to neutralize infection. These findings further mechanistic understanding of protein selectivity and may facilitate the development of diagnostics or therapeutics against henipavirus infection.

## INTRODUCTION

Nipah virus (NiV) was first identified following an outbreak in Malaysia in 1998-1999, where it was responsible for over a hundred deaths among people who were in close contact with farmed pigs and a million animals were culled to stop transmission (1, 2). Spillover events from certain Pteropus fruit bat species, which are a natural animal reservoir (3, 4), have continued in South and Southeast Asia. NiV can infect a broad range of mammalian species (5, 6). NiV infection has high morbidity and mortality in humans, and the virus is accordingly designated as a biosafety level-4 and category C priority pathogen. Due to its epidemic potential, the WHO has listed NiV as a priority disease for the research and development of therapeutics and vaccines (7).

NiV is an enveloped, negative-sense, single-stranded RNA paramyxovirus and a prototype species of the genus henipavirus. The NiV genome encodes six structural proteins. Two glycoproteins decorating the NiV surface orchestrate cell entry. The tetrameric attachment glycoprotein (G) binds to cell surface receptors ephrin-B2 (EFNB2) or ephrin-B3 (EFNB3) (8–10), triggering conformational changes in the trimeric fusion glycoprotein (F) resulting in the fusion of the viral and cell membranes (11, 12). NiV-G is a type II membrane glycoprotein with a short N-terminal cytoplasmic tail, followed by a transmembrane domain and a stalk domain that mediates tetrameric assembly and couples receptor-induced conformational changes to NiV-F fusion activity (12–17). The C-terminus of NiV-G is folded as a globular β-propeller domain that forms the terminal ‘head’ of the glycoprotein spike and is responsible for cell attachment (18–21). Multiple lines of evidence indicate that EFNB2 and EFNB3 are the cellular receptors (8–10, 17). Soluble EFNB2/3 has been shown to block NiV-F/G mediated membrane fusion and neutralize virus (8–10, 22), and soluble Eph receptors can also compete for EFNB2/3 on the host cell membrane to inhibit infection (22). NiV-G can bind EFNB2 and EFNB3 from diverse species (22), likely explaining the virus’ broad species tropism. Atomic resolution structures show that a loop (called the G-H binding loop) from EFNB2 makes extensive contacts to the central cavity of the β-propeller head domain, with notable contributions from hydrophobic side chains (19). The attachment glycoprotein (G) of Hendra virus (HeV), a close henipavirus relative, also binds EFNB2 and EFNB3 to mediate infection (8, 19, 22–24). The structures of EFNB2-bound NiV-G and HeV-G are almost identical with only a single conservative amino acid change at the interface (19).

Multiple NiV and HeV vaccine candidates have demonstrated protection in animal challenge models (5, 25, 26) by eliciting protective neutralizing antibody responses (5, 25–28). One subunit vaccine based on recombinant soluble HeV-G glycoprotein (HeV-sG) elicits cross-protective immune responses against both HeV and NiV; this vaccine is licensed and commercially developed for horses (29). Currently, the HeV-sG immunogen (26) and recombinant viral vector vaccine PHV02, which is a recombinant vesicular stomatitis virus (rVSV) expressing Ebola virus glycoprotein (EBOV-GP) and NiV-G, (30) are in phase 1 human clinical trials as part of the NiV vaccine candidates portfolio sponsored by the Coalition for Epidemic Preparedness Innovations (CEPI) (31). A mRNA candidate vaccine is also currently in phase 1 clinical trials (32). A human monoclonal antibody, m102.4, targeting both NiV-G and HeV-G has shown efficacy in animals (33–35). Since 2011, it has been administered as a post-exposure therapy on a compassionate basis for high-risk HeV cases in humans and has recently completed phase I clinical trials in Australia (36). The m102.4 antibody is also effective against a new HeV variant (37) that has been found circulating in fruit bats and horses in Australia (38–40). Generation of neutralization escape mutations is possible by virus passaging in the presence of low concentrations of antibody (41, 42). The use of antibody cocktails targeting non-competing neutralizing epitopes on G, or cocktails targeting both G and F glycoproteins, to restrict potential virus escape (21, 37, 43) is an area being tested.

Soluble receptors based on the cell entry receptors of viruses can be engineered as decoys with enhanced affinity (44) or specificity (45, 46) using deep mutational scanning (47). These decoys may be engineered to not participate in (i.e. are orthogonal to) the receptor’s normal biology (45, 46) and potently neutralize viruses *in vitro* (44–46, 48, 49). They have been shown to be broadly effective across diverse strains and variants (48, 50) and may perhaps be effective against related but yet to be identified viruses responsible for future zoonotic spillovers. Virus mutations for escape from soluble decoys have been found to be rare (50), and in principle, these mutations will also reduce binding to the native membrane-anchored receptor, thereby attenuating virulence or transmissibility. Since it was first used to identify EFNB2 as the NiV and HeV entry receptor (8, 9), soluble EFNB2 fused to the Fc-region of human immunoglobulin G1 (sEFNB2-Fc) has been shown to inhibit NiV and HeV fusion as well as fusion of closely related Cedar (51) (CedV) and Ghana (52) (GhV) henipaviruses *in vitro*, indicating soluble EFNB2 could be used for pan-specific treatment or prevention of henipaviral disease. Furthermore, sEFNB2-Fc inhibits more efficiently than the soluble EFNB3 Fc-fusion (10, 22, 51, 53). Additionally, soluble proteins are fused to Fc to enhance affinity through avidity, for improved serum stability, and to recruit immune effector functions (54). Only one closely-related henipavirus to date, Mòjiāng virus, does not engage EFNB2 to enter the cell (55). Despite its potential to be leveraged as an antiviral, EFNB2 is also an endogenous ligand for six ephrin (Eph) receptors: EphA4, EphB1 to EphB4, and EphB6 (56). These Eph-EFNB2 interactions regulate numerous important cell-cell signaling pathways (56), so wild-type sEFNB2-Fc may lead to off-target effects in humans. Studies in the past few years alone have utilized sEFNB2-Fc to explore its effects on cardiovascular disease (57–60), cancer (61–64), and other human diseases (65–67).

Crystal structures show that the interaction footprint of NiV-G and HeV-G on the EFNB2 surface (19) is nearly identical to that of Eph receptors (68–70), although atomic details and amino acid usage differ. In particular, there is a substantial shift in the position of the G-H binding loop (19). Here, we show how to use deep mutational scanning to impose specificity on the EFNB2 receptor binding domain (RBD) for binding NiV-G over Eph receptors. We then investigate how to engineer a sEFNB2-Fc variant that maintains virus interactions but has lost binding to Eph receptors, an idea previously proposed by Yuan *et al.* (71) and Xu *et al.* (72). We conclude that fine-tuning the conformational selectivity of the G-H binding loop is required to impose specificity on EFNB2 without a loss of affinity. The present study provides a foundation on how further optimization of sEFNB2-Fc could transform it into a potent antiviral countermeasure against NiV and henipaviral disease.

## RESULTS

### A deep mutational scan of EFNB2 using competition as a selection strategy

Previously, competition of soluble proteins for binding was employed as a selection strategy for deep mutagenesis of a membrane-bound receptor on the mammalian cell surface (45). By incubating the cells expressing receptor variants with the soluble proteins, the variants could be discriminated into populations that favor binding one soluble protein over the other (45). To achieve the same goal in this study, the head domain of NiV-G (a.a. G183 to T602) (73) was expressed as a soluble protein fused at the C-terminus with superfolder GFP (sfGFP) (74) in Expi293F cells, a suspension derivative of human HEK293 cells. Medium from soluble NiV-G-sfGFP expressing cells was incubated with Expi293F cells transfected with full-length EFNB2, and EFNB2 surface expression (via antibody staining of an extracellular N-terminal c-myc tag) and binding of the viral G glycoprotein were assessed by flow cytometry (**Fig. 1A**). NiV-G-sfGFP binding signal was proportional to EFNB2 expression, and the interaction was blocked by co-incubation with a representative Fc-fused Eph receptor to compete for EFNB2 binding sites (**Fig. 1B**). EphB2 was chosen because of its tight affinity for EFNB2 (75), and soluble EphB2-Fc (sEphB2-Fc) has been shown to inhibit infection of EFNB2 expressing cells by competing with NiV-G pseudoyped viruses (9).

**FIG 1.**
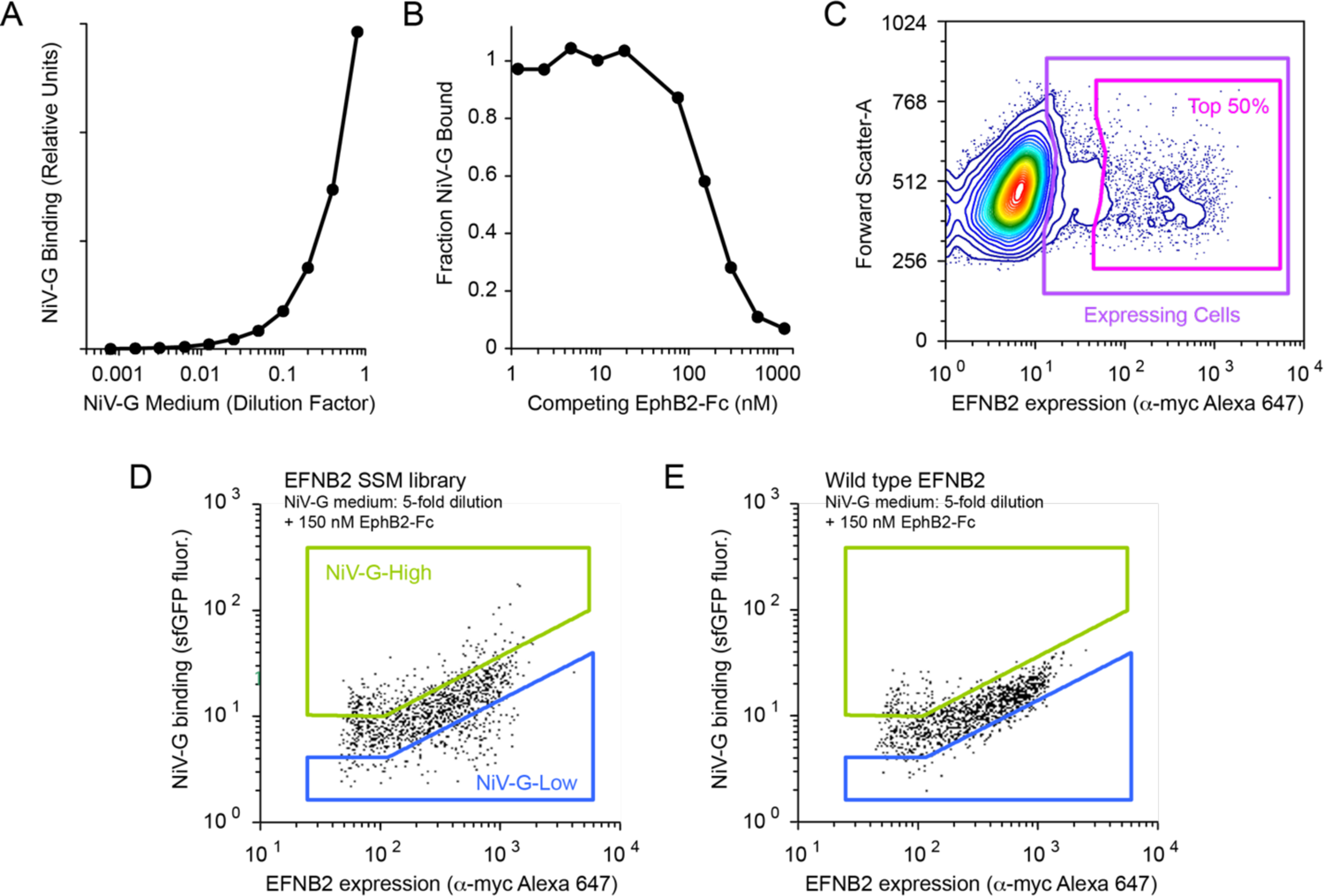
Competition as a selection strategy for identifying EFNB2 mutants specific for NiV-G. **(A)** EFNB2-expressing Expi293F cells were incubated with dilutions of NiV-G-sfGFP expression medium and bound NiV-G was detected by flow cytometry. **(B)** EFNB2-expressing cells were stained with a subsaturating 5-fold dilution of NiV-G-sfGFP medium. NiV-G binding was competed with increasing concentrations of sEphB2-Fc. **(C-D)** Expi293F cells transfected with an EFNB2 SSM library were co-incubated with a 5-fold dilution of NiV-G-sfGFP medium and 150 nM sEphB2-Fc. Under these conditions, EFNB2 mutants with either enhanced or decreased binding to NiV-G are resolved. After first gating viable cells based on scattering and propidium iodide exclusion, the top 50 % of EFNB2-expressing cells were gated (C, magenta). The upper and lower 15th percentiles for bound NiV-G-sfGFP fluorescence were collected in the NiV-G-High (D, green) and NiV-G-Low (D, blue) sorts. **(E)** Cells expressing wild-type EFNB2 analyzed under the same conditions. Note that the SSM library has EFNB2 variants with high and low NiV-G binding compared to wild-type.

A single-site saturation mutagenesis (SSM) library of EFNB2 was generated by overlap extension PCR encompassing all single amino acid substitutions in the extracellular RBD (a.a. I28 to G165, as numbered in PDB ID 2VSM and 2HLE). The library was then transfected into Expi293F cells under conditions that typically yield no more than a single coding variant per cell, linking cellular genotype to phenotype (76, 77). After incubation with a subsaturating 5-fold dilution of NiV-G-sfGFP medium in the presence of 150 nM sEphB2-Fc, the cells expressing the library were discriminated into three distinct populations based on NiV-G binding (**Fig. 1C and D**). High, wild-type-like, or low binding to NiV-G-sfGFP in the presence of competing sEphB2-Fc was delineated. In contrast, cells expressing wild-type EFNB2 exhibited only a single main population, which was indicative of no preferential binding to either soluble protein under the experimental conditions (**Fig. 1E**). Cells expressing EFNB2 variants with increased NiV-G binding specificity were expected to possess elevated sfGFP fluorescence due to diminished competition from EphB2. The collection of these cells by fluorescence-activated cell sorting (FACS) was referred to as the NiV-G-High sort **(green gate in Fig. 1D**). Within the same sorting experiment, cells expressing EFNB2 variants with diminished NiV-G-sfGFP binding, or possible enhanced binding to Eph receptors, were gated and collected, and this was referred to as the NiV-G-Low sort **(blue gate in Fig. 1D**). EFNB2 variants that fail to express at the plasma membrane were expected to be depleted from both sorted populations. Illumina sequencing was used to identify mutations with either high or low NiV-G binding in the sorted populations. By sequencing the naïve plasmid library and transcripts from the NiV-G-High and Low sorts, the enrichment ratios for all 2,760 amino acid substitutions in EFNB2-RBD were calculated (78). For the NiV-G-High sort, these ratios define a mutational landscape of EFNB2-RBD for enhanced NiV-G specificity **(Fig. S1)**. The data from two independent replicated experiments were correlated **(Fig. S2)**.

### A single mutation at the binding interface of EFNB2 increases pan-specificity to the attachment G glycoproteins of EFNB2-utilizing henipaviruses

Residue conservation scores for the NiV-G-High sort were calculated by averaging the log_2_ enrichment ratios for all substitutions at each residue in EFNB2-RBD. Mapping the conservation scores onto the crystal structures of EFNB2 bound to NiV-G (PDB ID 2VSM) (19) and EFNB2 bound to a human Eph receptor (EphB4, a close homolog of EphB2 with 79% similarity; PDB ID 2HLE) (69), shows that mutations that increase selectivity for NiV-G over EphB2 cluster to residues at the binding interface (**Fig. 2A**). In particular, substitutions at the base of the G-H binding loop are enriched; these substitutions may affect the conformation and dynamics of the loop. In contrast, the hydrophobic core of EFNB2-RBD is highly conserved as expected to maintain proper folding and expression. In the G-H binding loop, EFNB2-F120, L124, and W125 are relatively conserved due to their important interactions with hydrophobic pockets in NiV-G (18, 19). Site-directed mutagenesis studies have previously demonstrated that these residues are critical for NiV-G binding (10, 71) and their lower mutational tolerance in our scan further underscores their importance (**Fig. 2B and C**). EFNB2-F120, L124, and W125 also interact with hydrophobic surfaces in Eph receptors, and sites with overlapping or conserved functions for binding may not be useful in the context of imposing specificity. For instance, while a mutation to alanine at L124 has been shown to enhance NiV entry (71), it is poorly enriched in the NiV-G-High sort (**Fig. 2B**), indicating it has synchronous effects on binding to both the Eph receptor and NiV-G.

**FIG 2.**
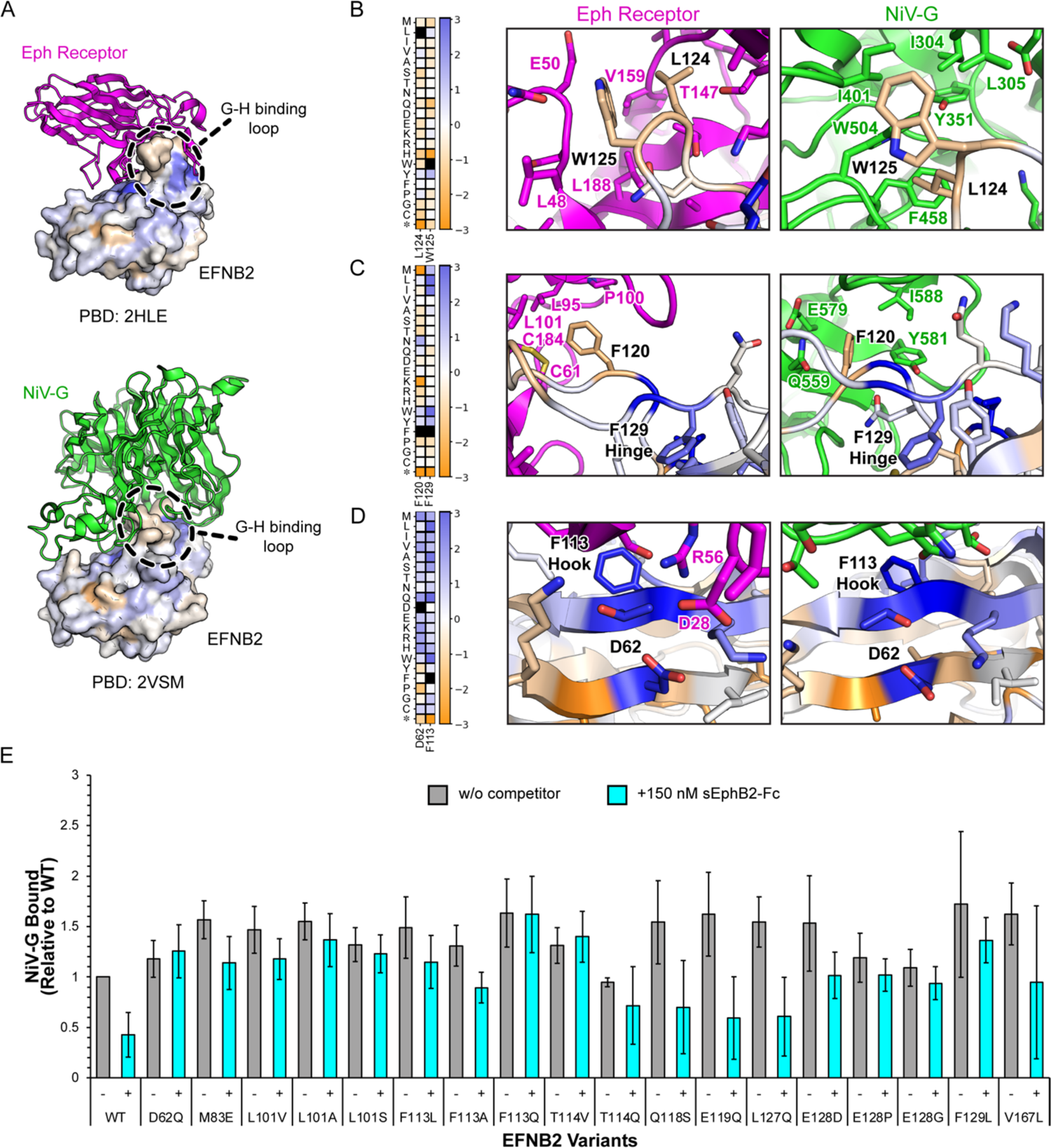
Mutations at the binding interface of EFNB2 enhance specificity to NiV-G over EphB2. **(A)** Conservation scores from the NiV-G-High sort are mapped onto the RBD of EFNB2 (shown as a surface) bound to Eph receptor (magenta) or NiV-G (green). For modeling EphB2, PBD ID 2HLE was chosen since EphB4 is a close homolog, sharing 79% similarity to EphB2. Residues in blue or orange indicate where mutations are enriched or depleted, respectively, in the mutational scan for preferential NiV-G binding. **(B-D)** Heat maps display the log_2_ enrichment ratios from the NiV-G-High sort, plotted from −3 (orange, depleted) to +3 (dark blue, enriched), for different interaction sites at the binding interface. Residue positions are on the horizontal axis, substitutions are on the vertical axis. Asterisks (*) denote stop codons. Wild-type amino acids are in black. Magnified views of interaction sites are shown on the right, with residues colored as in panel A. **(E)** Expi293F cells expressing wild-type or variants of myc-tagged EFNB2 were incubated with (cyan) or without (grey) sEphB2-Fc before the addition of a 5-fold dilution of NiV-G-sfGFP-medium. Binding of NiV-G was assessed by flow cytometry. Data are mean ± SD, N = 3 biological replicates.

Mutational “hotspots” for enhanced specificity may be located at residues with receptor-specific interactions, or at sites that do not interact directly with the binding partners but rather modulate the conformational dynamics of EFNB2. For example, EFNB2-F129 is located at the base of the G-H binding loop and is directed towards the core of EFNB2; it does not make any direct contacts with either NiV-G or Eph receptors (**Fig. 2C**). We hypothesize that EFNB2-F129 acts as a hinge for the G-H binding loop, and enriched substitutions of EFNB2-F129 identified in the mutational scan are proposed to shift the conformational equilibrium of the loop away from the ‘open’ state recognized by Eph receptors. In the ‘open’ state of the G-H binding loop bound by Eph receptors, EFNB2-L127 in the loop packs into a shallow hydrophobic pocket formed by EFNB2-M83, L101, F113, I115, and F129. In the ‘closed’ state bound by NiV-G, EFNB2-L127 shifts almost 5 Å out of the pocket as the G-H binding loop pivots around the F129 hinge. Enriched mutations that increase specificity for NiV-G include substitutions of EFNB2-F129 for leucine, isoleucine, and tryptophan; these mutations may favor the ‘closed’ G-H loop conformation while preserving interactions with the hydrophobic core required for folding and expression.

Other hotspots for specificity mutations are EFNB2 residues D62 and F113 that are found at the periphery and in the center of the binding interface, respectively. Many substitutions of these residues were enriched in the deep mutational scan for preferential binding to NiV-G. EFNB2-D62 participates in electrostatic interactions within a cluster of charged residues at the interface with an Eph receptor (**Fig. 2D**). This cluster is formed by an arginine and two glutamates in the EphB2 (E32, E45, and R57 from GenBank NM_004442.7) that are balanced by EFNB2-K60, D62, and K112 and is relatively conserved across all six Eph receptors (one glutamate is replaced by aspartate in EphB4 and is absent in EphB6) **(Fig. S3)**. No equivalent cluster of charged residues is found in the interface with NiV-G, and EFNB2-D62 makes no direct contacts with NiV-G. Many mutations to EFNB2-D62 are therefore selectively disruptive of Eph receptor binding. As a final example, EFNB2-F113 is proposed to act as a hook for high affinity binding to Eph receptors (**Fig. 2D**). When bound to an Eph receptor, EFNB2-F113 is rotated out of the hydrophobic core and oriented towards the binding interface where it forms a central hydrophobic anchor. However, when bound to NiV-G, the EFNB2-F113 hook is tucked back towards the EFNB2 hydrophobic core and only makes a glancing edge interaction with NiV-G. Many mutations to EFNB2-F113 were identified in the mutational scan as increasing specificity for NiV-G.

Based on their high enrichment in the NiV-G-High sort, eighteen mutations in EFNB2 were validated by targeted mutagenesis for enhanced NiV-G specificity (**Fig. 2E**). In the absence of competing soluble Eph receptor, the EFNB2 variants demonstrate close to if not greater than wild-type binding to soluble NiV-G. However, upon the addition of sEphB2-Fc competitor, most of the variants have small losses in NiV-G binding, whereas wild-type EFNB2 suffers from a large reduction. Although the deep mutational scan successfully identified EFNB2 mutants with preferential NiV-G binding, the mutations do not necessarily exhibit a complete loss of competition by EphB2. One mutation, D62Q, possesses high NiV-G binding that is unaffected by addition of sEphB2-Fc, demonstrating exceptional NiV-G specificity.

We sought to determine if a single mutation in EFNB2 (D62Q) affects its ability to bind to the attachment glycoproteins of closely related henipaviruses, since they share EFNB2 as an entry receptor (8, 79, 80) (**Fig. 3**). As with NiV-G, the head domains for HeV-G (a.a. G183 to S604) (24), GhV-G (a.a. T195 to Y632) (52), and CedV-G (a.a. K209 to C622) (51) were expressed as soluble proteins fused at the C-terminus with sfGFP in Expi293F cells. NiV-G shares 83% sequence identity with HeV-G (73), <30% with GhV-G (52), and ∼30% with CedV-G (79). The D62Q mutation in EFNB2 was chosen from the eighteen candidate mutations because, in addition to its NiV-G specificity, the D62Q mutation was predicted to not be affected by the sequence diversity among the viral G glycoproteins, as EFNB2-D62 does not make direct contacts with residues in NiV-G (**Fig. 2D**). For HeV-G and GhV-G, the D62Q mutant of EFNB2 demonstrated wild-type binding. For CedV-G, there was a slight loss of binding compared to wild-type.

**FIG 3.**
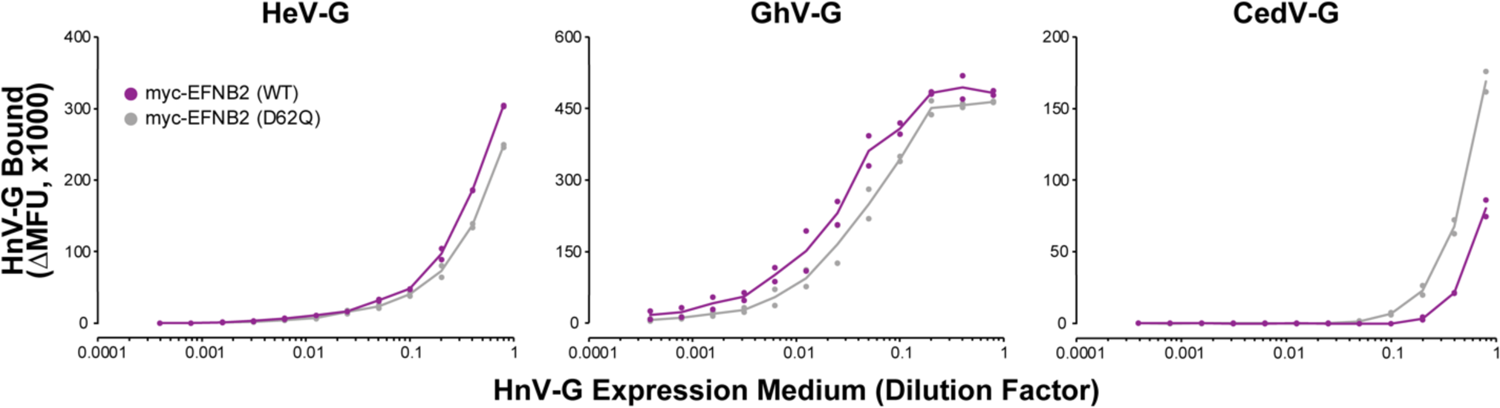
EFNB2-D62Q binds with pan-specificity to the attachment glycoproteins of henipaviruses (HnV-G). Expi293F cells expressing myc-tagged EFNB2, wild-type (grey) or the D62Q variant (purple), were incubated with dilutions of HnV-G-sfGFP medium. The three HnV-G tested were from Hendra virus (HeV-G, left), Ghana virus (GhV-G, middle), and Cedar virus (CedV-G, right). Bound HnV-G was detected by flow cytometry. Data points show range around the mean, N = 2 replicates.

### The D62Q variant of sEFNB2-Fc substantially reduces binding to four of its six Eph receptors

The six human Eph receptors (EphA4, EphB1 to EphB4, and EphB6) that endogenously interact with EFNB2 possess high structural homology but sequence similarity is dependent on the subclass (EphA4 versus EphB receptors), with differences primarily localized to the ephrin binding site (56). Crystal structures of EFNB2-Eph receptor complexes indicate that a conserved arginine on EphA4, EphB2, and EphB4 interacts with residue D62 on EFNB2 (68–70) **(Fig. S3)**. Assuming this interaction is common across all EFNB2-Eph receptor complexes, the D62Q mutation in EFNB2 may be sufficient to decrease affinity to the six Eph receptors. As mentioned previously, the potential for sEFNB2-Fc as an antiviral is hindered by possible off-target effects in humans due to Eph receptor binding. Therefore, for both wild-type and the D62Q variant, we tested the binding activity of sEFNB2-Fc against full-length Eph receptors expressed on the surface of transfected cells (**Fig. 4**). The sEFNB2-Fc construct fused the C-terminus of EFNB2-RBD to the N-terminus of the Fc region of human IgG1. The wild-type and D62Q variant proteins were then purified through affinity capture followed by size exclusion chromatography (SEC) for high purity **(Fig. S4)**. Using flow cytometry to measure binding of the purified sEFNB2-Fc proteins, the D62Q mutation was shown to reduce binding to all six Eph receptors expressed at the cell surface when compared to wild-type. The most pronounced decreases were observed for EphA4, EphB1, EphB2, and EphB6. The D62Q variant exhibited high residual binding to EphB3 and EphB4, possibly because of additional high affinity interactions not conserved among all EFNB2-Eph receptor complexes.

**FIG 4.**
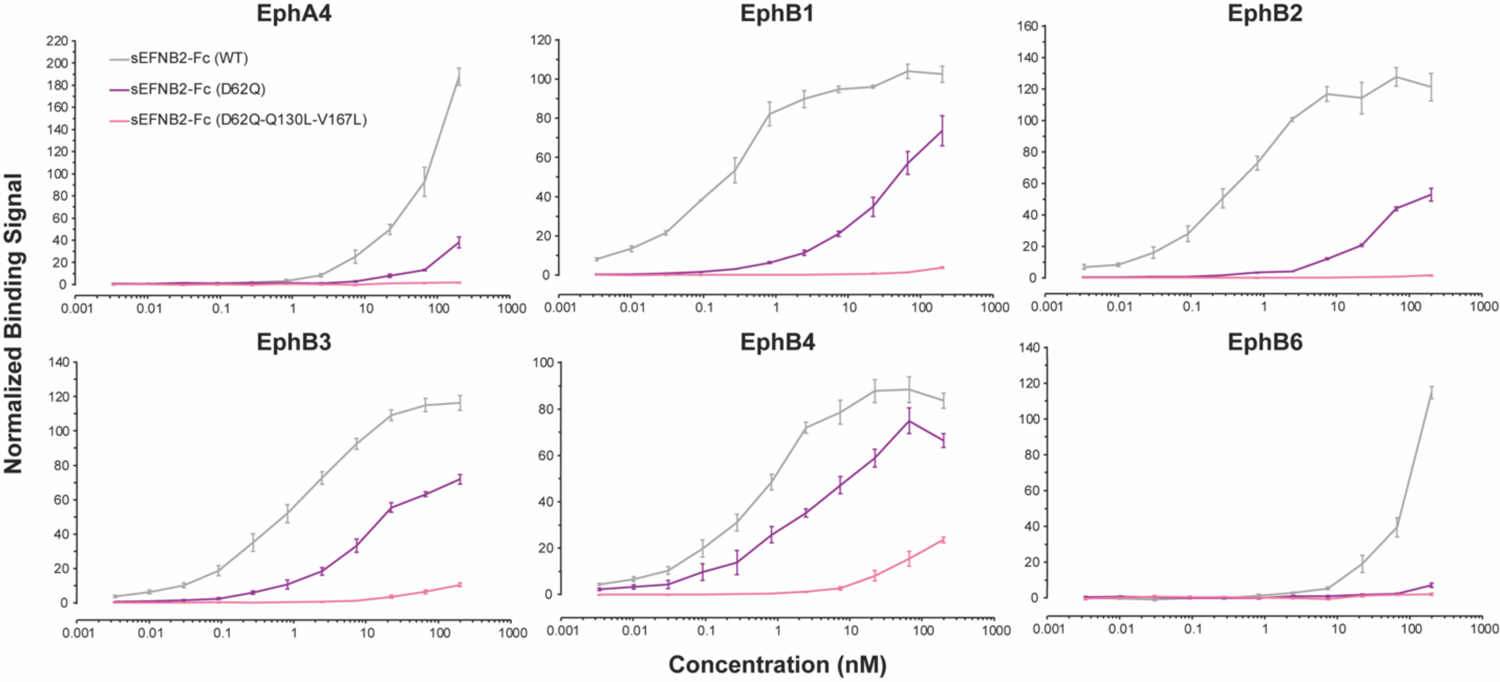
Variants of sEFNB2-Fc with decreased binding to Eph receptors. Expi293F cells were transfected with expression plasmids for full-length myc-tagged Eph receptors. The six Eph receptors that bind EFNB2 in humans are EphA4 (top left), EphB1 (top middle), EphB2 (top right), EphB3 (bottom left), EphB4 (bottom middle), and EphB6 (bottom right). Binding of sEFNB2-Fc to surface-expressed Eph receptors was measured by flow cytometry. Data are mean ± SEM, N = 3 biological replicates.

### Higher order combinations of mutations in EFNB2 further increase specificity towards NiV-G

The deep mutational scan of EFNB2-RBD in the wild-type background identified many candidate mutations that could be combined with the D62Q mutation to further improve NiV-G specificity. However, it is ambiguous how two mutations in EFNB2-RBD might interact based on rational inspection of available structures. Two major considerations for combining mutations to reduce Eph receptor binding are: (1) the mutations may also diminish binding to NiV-G, and (2) the mutations may affect the stability of sEFNB2-Fc. Both are difficult to predict from the deep mutational scan of full-length EFNB2. Nevertheless, we hypothesized that a second deep mutational scan of EFNB2-RBD in the D62Q background, this time using competition between NiV-G-sfGFP and soluble Fc-fusions of EphB3 and EphB4 (the two Eph receptors that have persistent binding to sEFNB2-Fc-D62Q), may reveal secondary substitutions for enhanced NiV-G specificity. We generated a SSM library of EFNB2-RBD with the D62Q mutation fixed. The library of sequences was transfected into Expi293F cells using the same conditions as the deep mutational scan in the wild-type background. For selection by FACS, the cells expressing the EFNB2-D62Q library were incubated with a subsaturating 5-fold dilution of NiV-G-sfGFP medium in the presence of equimolar concentrations of sEphB3-Fc and sEphB4-Fc (1 μM of each). The NiV-G-High and Low gated populations were again collected for Illumina sequencing to identify mutations with either high or low NiV-G binding in the presence of the competitors. The enrichment ratios for every single-site substitution and the conservation score at each position were calculated as described for the scan in the wild-type EFNB2 background **(Fig. S5)**. The data from two independent replicates were highly correlated **(Fig. S6)**.

The difference was calculated between the conservation scores for the EFNB2 deep mutational scans on the wild-type versus D62Q backgrounds. Once mapped onto the crystal structures of EFNB2 bound to NiV-G (PDB ID 2VSM) (19) or EphB4 (PDB ID 2HLE) (69), the difference scores highlight that by fixing the D62Q substitution, EFNB2 has become markedly less tolerant of mutations for maintaining high NiV-G binding in the presence of competing Eph receptors (**Fig. 5A; Fig. S7)**. This is consistent with the EFNB2-D62Q sequence being more optimized for tight and specific binding to NiV-G, with most additional mutations being deleterious **(Fig. S7A)**. Higher sequence conservation in the EFNB2-D62Q background is especially apparent in the hydrophobic core and at the G-H binding loop. The second deep mutational scan confirmed predictions that some candidate mutations from the first scan are not tolerated in combination with D62Q, based on their low enrichment. On the other hand, the second scan did identify some highly enriched substitutions, both new and shared with the first scan, as possible candidates for combining with D62Q. Eight EFNB2 double mutants were validated by targeted mutagenesis for enhanced NiV-G binding in the presence of competing sEphB3-Fc and sEphB4-Fc (**Fig. 5B**). However, there remained partial competition by soluble Eph receptors for NiV-G binding for all of the EFNB2 double mutants. Furthermore, several of the EFNB2 double mutants demonstrated lower binding to NiV-G in the absence of any competitor, suggesting decreased affinity for the viral G glycoprotein.

**FIG 5.**
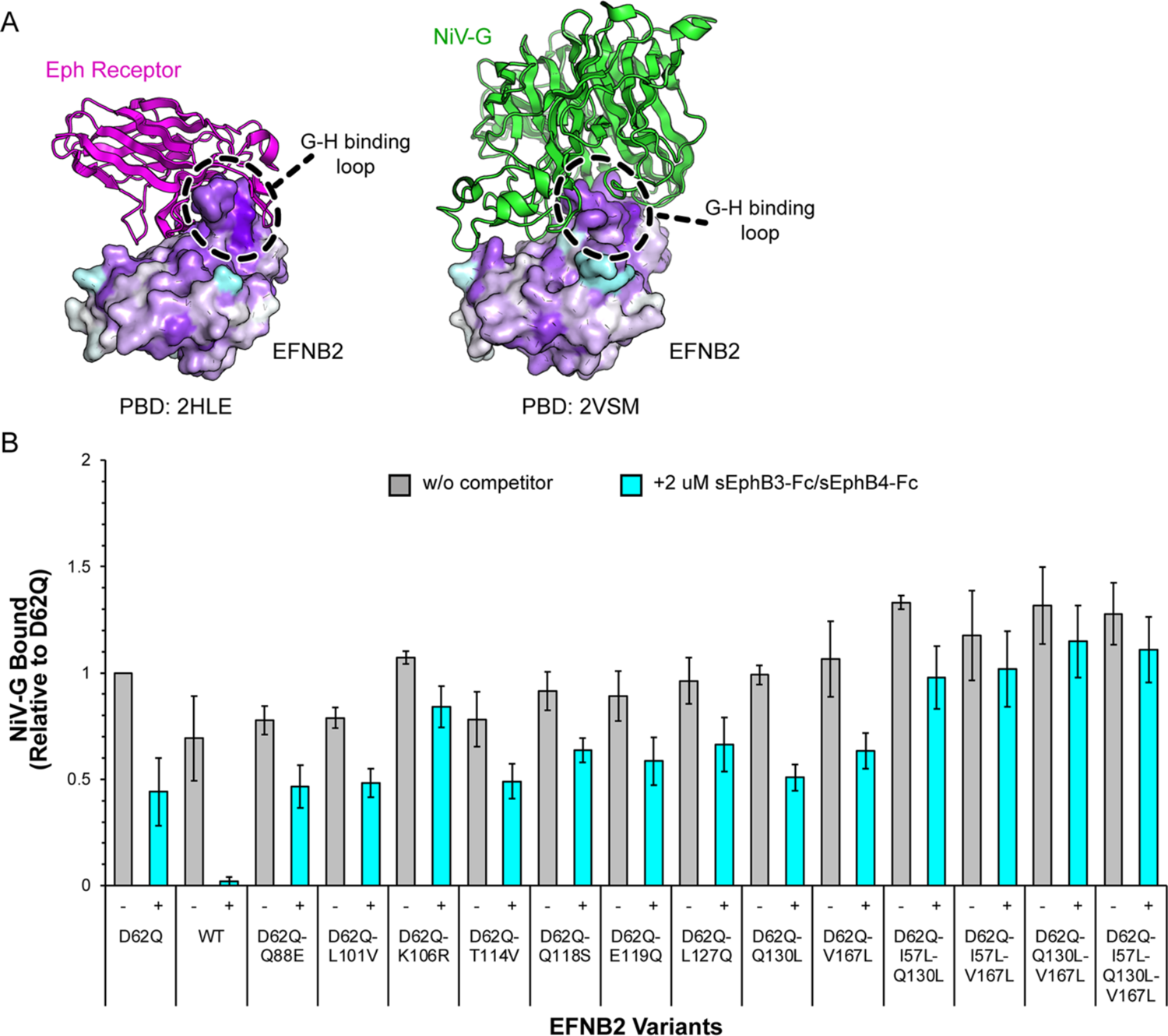
A deep mutational scan for NiV-G specificity of EFNB2 in the D62Q background reveals reduced mutational tolerance. **(A)** The difference of the conservation scores for the NiV-G-High sorts from the two EFNB2 deep mutational scans are mapped onto the RBD of EFNB2 bound to Eph receptor (magenta) or NiV-G (green). Residues in cyan are more conserved in the deep mutational scan of wild-type EFNB2, while those in dark purple are more conserved in the deep mutational scan of EFNB2-D62Q. **(B)** Expi293F cells expressing wild-type or variants of myc-tagged EFNB2 were incubated with (cyan) or without (grey) equimolar amounts (1 μM) of sEphB3-Fc and sEphB4-Fc before the addition of a 5-fold dilution of NiV-G-sfGFP medium. Binding of NiV-G was assessed by flow cytometry. Data are mean ± SD, N = 3 biological replicates.

To address the issue that the EFNB2 double mutants remained partially selective for NiV-G, we next tested combinations of three mutations chosen from the set of I57L, Q130L, and V167L, plus D62Q. These were chosen based on their high enrichment in the deep mutational scan of EFNB2-D62Q and in the case of Q130L and V167L, their high NiV-G binding in the absence of competitor (**Fig. 5B**). As with the EFNB2 double mutants, three triple mutants and one quadruple mutant were validated by targeted mutagenesis of full-length EFNB2 (**Fig. 5B**). All four higher order mutants of EFNB2 had greater binding to NiV-G in the presence of competitor than the EFNB2-D62Q single mutant. Soluble EFNB2-Fc carrying the D62Q-Q130L-V167L mutations was purified through affinity capture and SEC **(Fig. S4)**. Dilutions of the purified sEFNB2-Fc triple mutant were then incubated with Expi293F cells expressing full-length Eph receptors and the binding was quantified using flow cytometry (**Fig. 4**). The three substitutions in sEFNB2-Fc led to substantially diminished binding to the panel of six human Eph receptors, in contrast to the binding activity of wild-type sEFNB2-Fc (binds all six receptors tightly) and the D62Q variant (binds EphB3 and EphB4 moderately).

### The D62Q-Q130L-V167L variant of sEFNB2-Fc has reduced binding to NiV-G

In a previous study, a single mutation was sufficient to confer orthogonality to a soluble virus receptor, meaning it no longer participates in binding interactions with endogenous ligands, yet the receptor maintained wild-type binding to the virus spike (45). Three substitutions were required to knock out endogenous Eph interactions in sEFNB2-Fc, but it was unknown if the substitutions affected affinity for NiV-G. To test this, binding of wild-type and variants of sEFNB2-Fc to full-length NiV-G expressed on the surface of Expi293F cells was measured via flow cytometry (**Fig. 6B**). The sEFNB2-Fc-D62Q variant bound to NiV-G similarly to wild-type sEFNB2-Fc, but the addition of the Q130L and V167L mutations reduced binding to the viral G glycoprotein. These observed binding activities were further confirmed by measuring the neutralization activities of the sEFNB2-Fc proteins against replication-competent recombinant Cedar virus (rCedV) expressing a GFP reporter (rCedV-GFP) and chimeric rCedV-GFP whereby the G and F glycoproteins of CedV were replaced with either the NiV Bangladesh strain (NiV_B_) or HeV G and F glycoproteins (rCedV-NiV_B_-GFP and rCedV-HeV-GFP, respectively). This system was previously established as a biosafety level-2 (BSL-2) alternative for NiV/HeV neutralization assays (21, 37, 43). For rCedV-NiV_B_-GFP and rCedV-HeV-GFP, the D62Q variant of sEFNB2-Fc demonstrated wild-type-like neutralizing activity with comparable IC_50_ values (approximately 20-50 pM) between the two soluble decoys (**Fig. 6C) (Table 1**). This wild-type-like neutralization activity of the sEFNB2-Fc-D62Q variant confirmed what was observed previously in this study for binding to NiV-G (**Fig. 6B**) and HeV-G (**Fig. 3**). Furthermore, the D62Q variant of sEFNB2-Fc exhibited lower neutralizing activity against rCedV-GFP virus compared to wild-type sEFNB2-Fc, which correlated well with the loss of binding observed previously in this study against CedV-G (**Fig. 3**). The triple mutant of sEFNB2-Fc, on the other hand, had diminished neutralization activity against all three chimeric viruses and possessed higher IC_50_ values (in the nanomolar range) in contrast to those of wild-type and the D62Q variant of sEFNB2-Fc (**Table 1**).

**FIG 6.**
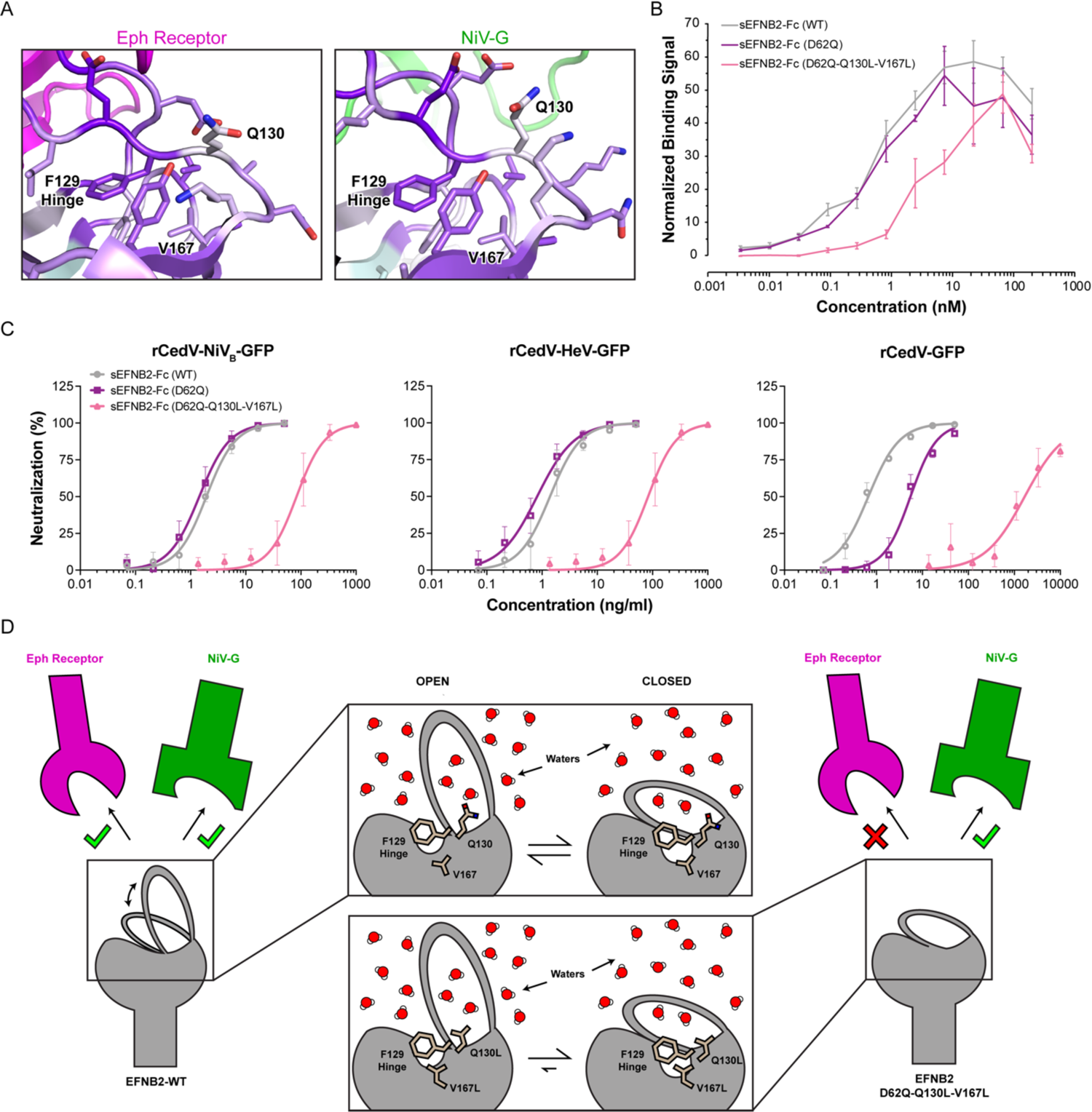
Mutations at the base of the G-H binding loop increase specificity of EFNB2 but at the cost of decreased NiV-G binding. **(A)** Magnified views of the G-H binding loop bound to Eph receptor (magenta) and NiV-G (green) are shown. The difference of the conservation scores for the NiV-G-High sorts from the two EFNB2 deep mutational scans are mapped onto the RBD of EFNB2 and colored as in Figure 5A. **(B)** Expi293F cells expressing full-length NiV-G fused at the C-terminus to sfGFP were incubated with increasing concentrations of sEFNB2-Fc protein. Binding of sEFNB2-Fc to membrane-displayed NiV-G-sfGFP was measured by flow cytometry. Data are mean ± SEM, N = 3 biological replicates. **(C)** Mixtures of rCedV-GFP chimeric viruses pre-incubated with increasing concentrations of sEFNB2-Fc protein were added to pre-seeded Vero 76 cells. Neutralizing activity of sEFNB2-Fc was measured by counting the reduction in GFP fluorescent foci. Data are mean ± SD, N = 2 independent experiments, each performed in technical triplicate. **(D)** Proposed mechanism by which the Q130L and V167L mutations affect the conformational selectivity of the G-H binding loop. In wild-type EFNB2, the G-H binding loop is dynamic and in equilibrium between conformations for binding both Eph receptors (magenta) and NiV-G (green). The side chain of EFNB2-Q130 is fully exposed to solvent in the ‘open’ loop structure bound to Eph receptors but is packed against neighboring side chains in the ‘closed’ loop structure bound by NiV-G. In EFNB2-D62Q-Q130L-V167L, the conformational equilibrium shifts to favor the ‘closed’ state, which minimizes interactions between solvent and the hydrophobic Q130L substitution. Additionally, a leucine substitution at EFNB2-V167 acts as a cavity-filling mutation and increases the hydrophobic packing around EFNB2-F129 in the ‘closed’ state.

**TABLE 1.** Neutralizing Activity of Soluble Decoy Receptors

| Chimeric Virus | sEFNB2-Fc |  |  |
| --- | --- | --- | --- |
|  | Wildtype | D62Q | D62Q-Q130L-V167L |
|  | IC<sub>50</sub> ng/ml<br>(nM) |  |  |
| rCedV-NiV<sub>B</sub>-GFP | 1.93 ± 0.18<br>(0.0466 ± 0.0043) | 1.44 ± 0.17<br>(0.0348 ± 0.0041) | 83 ± 14<br>(2.01 ± 0.35) |
| rCedV-HeV-GFP | 1.39 ± 0.24<br>(0.0336 ± 0.0058) | 0.81 ± 0.13<br>(0.0196 ± 0.0031) | 35.8 ± 4.8<br>(0.87 ± 0.12) |
| rCedV-GFP | 0.67 ± 0.07<br>(0.0162 ± 0.0017) | 5.49 ± 0.71<br>(0.133 ± 0.017) | 1,680 ± 480<br>(41 ± 11) |

In the crystal structure with NiV-G, residue EFNB2-V167 is buried in the protein core immediately below EFNB2-F129 (the ‘hinge’) and does not interact directly with NiV-G, whereas residue Q130 is at the base of the G-H binding loop and is in close enough proximity to participate in hydrogen bonding with NiV-G, although no hydrogen bonds are observed in the static model built into the crystallographic electron density (19) (**Fig. 6A**). An alanine substitution at EFNB2-Q130 increases infectivity by NiV-G pseudotyped virus, suggesting that any hydrogen bonds EFNB2-Q130 might make are not critical for NiV-G binding (71). We hypothesize instead that the Q130L and V167L mutations modulate the conformation of the G-H loop for selective recognition and binding of receptors (**Fig. 6D**). The side chain of EFNB2-Q130 extends towards solvent in Eph receptor-bound structures but in the NiV-G-bound ‘closed’ state, EFNB2-Q130 is surrounded on its sides by other residues forming the base of the G-H loop (19, 68–70); the percent of solvent accessible surface area for EFNB2-Q130 is 59.5% when an Eph receptor is bound (calculated from PDB 2HLE), decreasing to 30.0% when NiV-G is bound (calculated from PDB 2VSM) (81). A hydrophobic leucine substitution at this position is expected to favor the ‘closed’ state of the G-H loop to minimize exposure to solvent. Likewise, the leucine substitution at EFNB2-V167 is speculated to readily accommodate the ‘closed’ conformation recognized by NiV-G by acting as a cavity-filling mutation and increasing hydrophobic packing around the G-H loop hinge (EFNB2-F129). We cannot exclude the possibility that other higher order combinations of substitutions may better fine-tune the conformational dynamics and surface properties of EFNB2 to enhance NiV-G specificity without a loss in affinity.

## DISCUSSION

The power of deep mutational scanning for designing selective ligands lies within the ability to identify single substitutions that alter a protein’s specificity. Here, a deep mutational scan of the RBD of EFNB2 highlighted how the binding interface, especially at the base of the G-H binding loop, contained hotspots for mutations that enhance specificity for NiV-G over Eph receptors. Mutations at these sites were hypothesized to either influence the conformational dynamics of the G-H binding loop, thus altering the equilibrium between a ‘closed’ and ‘open’ state recognized by NiV-G or Eph receptors, respectively, or alternatively remove key interactions for Eph receptor binding but not for NiV-G binding. Indeed, the D62Q variant of EFNB2 almost eliminated binding to four of six human Eph receptors and diminished binding to the remaining two Eph receptors, while preserving pan-specific binding to G of four closely related henipaviruses (NiV, HeV, GhV, and CedV) that all use EFNB2 as a cell entry receptor. The sEFNB2-D62Q-Fc variant potently neutralized the human henipaviruses NiV and HeV. The mutational landscape of EFNB2 also highlighted other residues as important specificity determinants, including a phenylalanine hook (F113) that rotates away from the EFNB2 core to engage Eph receptors, and a phenylalanine hinge (F129) that acts as the pivot point at which the G-H loop bends.

While the specificity of sEFNB2-Fc towards NiV-G is greatly enhanced through a single D62Q mutation, it is unclear whether residual binding to EphB3 and EphB4 will impart unacceptable toxicity or lead to target-mediated disposition *in vivo*. We therefore sought to increase specificity further by combining multiple mutations, but this is challenging from a deep mutational scan when only single substitutions are sampled in the library. Epistatic interactions between mutations are often ambiguous and the effects of combining mutations become difficult to predict. We chose to fix the D62Q mutation in EFNB2 and perform a second deep mutational scan for NiV-G specificity. In this second scan, the mutational landscape of EFNB2 became less tolerant of mutations, consistent with the starting sequence already being partially optimized for tight and specific binding to NiV-G. There were also epistatic interactions that shifted amino acid preferences at multiple sites when EFNB2 was scanned in the D62Q versus wild-type background. Rational combinations of mutations based on this second scan produced a highly specific soluble EFNB2 for henipaviruses. This triple mutant of EFNB2 showed negligible residual binding to Eph receptors, but it also suffered from reduced potency (low nanomolar versus picomolar IC_50_ values) against NiV, HeV, and CedV in a chimeric virus neutralization assay. Our results emphasize that designing ligands for enhanced specificity, without a loss of affinity, is exceedingly challenging when there are multiple interaction partners that share an almost identical binding surface. One avenue of exploration is the use of machine learning prediction tools trained on deep mutagenesis data sets to computationally design the best combinatorial mutations.

Soluble decoy receptors with high affinity and specificity for the EFNB2-utilizing henipaviral G glycoproteins may have uses not only as therapeutics but also as diagnostics. Although the D62Q variant of sEFNB2-Fc has residual binding to Eph receptors that warrants caution for use as a therapy, it also has potential as a useful diagnostic for the detection of henipaviral infection. Currently, the two primary methods of virus detection are antibody-based methods (e.g. enzyme-linked immunoassay for henipaviral proteins) or polymerase chain reaction amplification of henipaviral genes (6). Mutation or unknown variants of known henipaviruses, or a future zoonotic spillover of a novel EFNB2-utilizing henipavirus, may render antibodies and/or diagnostic PCR primers useless and require the production of new diagnostics, but so long as the henipaviral G glycoprotein engages EFNB2 as an entry receptor, sEFNB2-Fc and likely sEFNB2-D62Q-Fc will continue to detect such viruses. Future deep mutational scanning of henipaviral G may also help to understand the sequence space available for evasion of diagnostics as well as therapeutics and may illuminate the pandemic potential of these deadly viruses.

## METHODS

### Plasmids

Human EFNB2 isoform 1 (GenBank NP_004084.1) was cloned as the mature polypeptide (amino acids (a.a.) I28 to V333) into the NheI-XhoI sites of pCEP4 (Invitrogen) with a N-terminal HA leader (MKTIIALSYIFCLVFA), c-myc epitope tag, and linker (GSPGGA). Soluble EFNB2-Fc was constructed by cloning the receptor binding domain (a.a. I28 to G165) of EFNB2 into the NheI-BamHI sites of pcDNA3.1(+) (Invitrogen) with a N-terminal CD5 leader (MPMGSLQPLATLYLLGMLVASVLA) and a C-terminal linker (GS) followed by the Fc region of human IgG1 (GenBank KY432415.1; a.a. D221 to K447). Synthetic human codon-optimized gene fragments (Integrated DNA Technologies) for the head domains of NiV-G (GenBank NP_112027.1, a.a. G183 to T602), HeV-G (GenBank NP_047112, a.a. G183 to S604), GhV-G (GenBank AFH96011.1, a.a. T195 to Y632), and CedV-G (GenBank AFP87279.1, a.a. K209 to C622) were cloned into the NheI-BamHI sites of pcDNA3.1(+), each with a N-terminal CD5 leader and a C-terminal linker (GSGGSGSGG) followed by superfolder GFP (sfGFP) (GenBank ASL68970) (74). Full-length NiV-G fused with a C-terminal sfGFP was constructed by using overlap extension PCR to assemble the synthetic human codon-optimized gene fragments (Integrated DNA Technologies) for the N-terminus of NiV-G (a.a. M1 to V182) and the C-terminal head domain (a.a. G183 to T602) into a single gene fragment. The resulting assembled fragment was cloned into the NheI-BamHI sites of pcDNA3.1(+) with a C-terminal linker (GSGGSGSGG) followed by sfGFP. Human codon-optimized full-length Eph receptors were subcloned from pDONR223-EphB1 (Addgene # 23930 (82)), pMD18-T-EphB2 (Sino Biological), pMD18-T-EphB4 (Sino Biological), and pMD18-T-EphB6 (Sino Biological) into the NheI-XhoI sites of pCEP4 (Invitrogen) with a N-terminal HA leader, c-myc tag, and linker (GSPGGA). For soluble EphB3-Fc and EphB4-Fc, the extracellular domains of EphB3 and EphB4 were subcloned from pCMV3-SP-N-Myc-EphB3 (Sino Biological) and pCEP4-myc-EphB4, respectively, into the NheI-BamHI sites of pcDNA3.1(+) (Invitrogen) with a N-terminal CD5 leader and a C-terminal linker (GS) followed by the Fc region of human IgG1 (a.a. D221 to K447).

### Protein Expression in Expi293F Tissue Culture

Expi293F cells (ThermoFisher Scientific) were cultured in Expi293 Expression Medium (ThermoFisher Scientific) at 125 rpm, 8% CO_2_, and 37°C. Cells were transfected at a density of 2 × 10^6^ cells/mL. For each mL of culture, 500 ng of plasmid was mixed with 3 μg of polyethylenimine (MW 25,000; Polysciences) or Expifectamine (ThermoFisher Scientific) in 100 μL of OptiMEM (Gibco), incubated for 20 minutes at room temperature, and added to cells. Transfection Enhancers (ThermoFisher Scientific) were added 18-23 h later if using polyethylenimine or 16-18 h later if using Expifectamine. Transfected cells were cultured for 5 days before harvesting the medium supernatant. Cells were removed by centrifugation at 800 x g for 10 minutes at 4°C, and then cell debris and precipitates were removed by centrifugation at 24,000 x g for 20 minutes at 4°C. For soluble sfGFP fusions of henipaviral G head domains (NiV-G-sfGFP, HeV-G-sfGFP, CedV-G-sfGFP, GhV-G-sfGFP), the clarified expression medium was stored at −20°C and not purified further.

### Purification of IgG1-Fc Fused Proteins

Clarified expression medium was incubated with KANEKA KanCapA 3G Affinity resin (Pall; equilibrated in phosphate buffered saline (PBS)) for 90 minutes at 4°C. Resin was collected on a chromatography column and washed with 10 column volumes (CV) PBS. The protein was eluted with 5 CV 60 mM sodium acetate pH 3.7, and 1 CV 1 M Tris pH 9.0 was added to the eluate to neutralize the pH. The eluted proteins were concentrated with a centrifugal device (MWCO 30 kDa for soluble EFNB2 proteins and 50 kDa for soluble Eph receptor proteins; Sartorius) before being separated on a Superdex 200 Increase 10/300 GL (Cytiva Life Sciences) with PBS as the equilibration and running buffers. Peak fractions were pooled, concentrated, and stored at −80°C after snap freezing in liquid nitrogen. The protein concentrations were determined by absorbance at 280 nm using the calculated molar extinction coefficients for the monomeric mature polypeptide sequences.

### Deep Mutagenesis

A SSM library focused on the RBD of EFNB2 (a.a. I31 to G168) was constructed using pCEP4-myc-EFNB2 as the template, which encodes myc-tagged, full-length EFNB2. In the second scan, the SSM library contained a fixed D62Q mutation, while all other positions were mutagenized. Overlap extension PCR was used to introduce degenerate NNK codons at every position in EFNB2-RBD (83). The library was transfected into Expi293F cells under conditions that typically yield no more than a single coding variant per cell (76, 77). Cells were transfected with 1 ng of coding plasmid diluted in 1.5 μg of pCEP4-△CMV carrier plasmid (77) per mL of culture at 2 × 10^6^ cells/mL using Expifectamine (ThermoFisher Scientific). The medium was replaced 2 h post-transfection. At 24 h post-transfection, the cells were harvested for fluorescence-activated cell sorting (FACS). Cells were washed with ice-cold PBS supplemented with 0.2% w/v bovine serum albumin (PBS-BSA). Cells were then incubated for 5 minutes on ice with a mixture of soluble human IgG1-Fc fused Eph receptor/s (150 nM sEphB2-Fc (R&D systems) for the first scan or 1 μM sEphB3-Fc plus 1μM sEphB4-Fc for the second scan) and anti-myc Alexa 647 (clone 9B11, 1/250 dilution, Cell Signaling Technology) in PBS-BSA before the addition of NiV-G-sfGFP-containing medium (1/5 dilution). The mixture was incubated on ice for 30 minutes, washed twice with PBS-BSA, and sorted on a BD FACS Aria II at Roy J. Carver Biotechnology Center. The main cell populations were gated by forward/side scattering to remove debris and doublets, while propidium iodide staining was used to exclude dead cells. From the myc-positive (Alexa 647) population, the 10% of cells with the highest and the 15% with the lowest GFP fluorescence were collected (**Fig. 1D**). The collection tubes were prepared by coating overnight with fetal bovine serum, and Expi293 Expression Medium was added to the tubes prior to sorting. Total RNA was extracted from the collected cells using a GeneJET RNA purification kit (ThermoFisher Scientific). Complementary DNA (cDNA) was reverse transcribed with Accuscript (Agilent) primed with a gene-specific oligonucleotide. The mutagenized region of EFNB2 was PCR amplified as two overlapping fragments. A second round of PCR was performed on the fragments to add adapters for unique barcoding, annealing to Illumina sequencing primers, and binding to the flow cell. The PCR products were sequenced on an Illumina NovaSeq 6000 using 2 × 250 nucleotide paired end protocol, and the resulting data were analyzed using Enrich (78). The frequencies of EFNB2 variants in the transcripts of sorted populations were compared to their frequencies in the naïve plasmid library. The log_2_ enrichment ratios for each variant were calculated and normalized by subtracting the log_2_ enrichment ratio for the wildtype sequence. Conservation scores at each residue position were calculated by averaging the log_2_ enrichment ratios for all nonsynonymous mutations at the same position.

### Competition Assay

Expi293F cells were transfected with 500 ng of pCEP4-myc-EFNB2 (wildtype or mutants) per mL of culture at 2 × 10^6^ cells/mL using Expifectamine (ThermoFisher Scientific). At 24 h post-transfection, the cells were harvested for analysis. A competitor solution was prepared by mixing soluble human IgG1-Fc fused Eph receptor/s (150 nM sEphB2-Fc or 1 μM sEphB3-Fc plus 1μM sEphB4-Fc) and anti-myc Alexa 647 (clone 9B11, 1/250 dilution, Cell Signaling Technology) in PBS-BSA. A non-competitor solution was prepared by diluting anti-myc Alexa 647 (clone 9B11, 1/250 dilution, Cell Signaling Technology) in PBS-BSA. Transfected cells were washed with PBS-BSA and incubated with non-competitor or competitor solution for 5 minutes on ice before the addition of NiV-G-sfGFP-containing medium (1/5 dilution). The mixture was incubated on ice for 30 minutes, washed twice with PBS-BSA, and analyzed on a BD Accuri C6 flow cytometer.

### Henipaviral G Head Domain Titration Assay

Expi293F cells were transfected with 500 ng of pCEP4-myc-EFNB2 (wildtype or mutants) per mL of culture at 2 × 10^6^ cells/mL using Expifectamine (ThermoFisher Scientific). At 24 h post-transfection, the cells were harvested for analysis. The cells were washed with PBS-BSA and incubated with one-half serial dilutions of expression medium containing HeV-G-sfGFP, GhV-G-sfGFP, or CedV-G-sfGFP for 20 minutes at 4°C. After washing once with PBS-BSA, the cells were incubated with anti-myc Alexa 647 (clone 9B11, 1/250 dilution, Cell Signaling Technology) in PBS-BSA for 20 minutes at 4°C. Cells were washed twice with PBS-BSA and analyzed on a BD Accuri C6 flow cytometer.

### Eph Receptor Binding Assay

Expi293F cells were transfected with 500 ng of plasmid expressing myc-tagged full-length Eph receptor (pCEP4-myc-Eph for EphB1, B2, B4, and B6; pCMV3-SP-N-Myc-EphA4 and pCMV3-SP-N-Myc-EphB3, Sino Biological) per mL of culture at 2 × 10^6^ cells/mL using Expifectamine (ThermoFisher Scientific). At 24 h post-transfection, the cells were harvested for analysis. The cells were washed with PBS-BSA and incubated with one-third serial dilutions of sEFNB2-Fc for 20 minutes at 4°C. After washing once with PBS-BSA, the cells were incubated with anti-myc-fluorescein isothiocyanate (FITC) (chicken polyclonal, 1/150 dilution, Immunology Consultants Laboratory) and anti-human IgG-allophycocyanin (APC) (clone M1310G05, 1/250 dilution, BioLegend) in PBS-BSA for 20 minutes at 4°C. Cells were washed twice with PBS-BSA and analyzed on a BD Accuri C6 flow cytometer. Binding signal was calculated by subtracting background fluorescence of cells incubated without sEFNB2-Fc and normalized across replicates based on total measured fluorescence for each day of experiments.

### NiV-G Binding Assay

Expi293F cells were transfected with 500 ng of pcDNA3-NiV-G-sfGFP (full-length) per mL of culture at 2 × 10^6^ cells/mL using Expifectamine (ThermoFisher Scientific). At 24 h post-transfection, the cells were harvested for analysis. The cells were washed with PBS-BSA and incubated with one-third serial dilutions of sEFNB2-Fc for 20 minutes at 4°C. After washing once with PBS-BSA, the cells were incubated with anti-human IgG-APC (clone M1310G05, 1/250 dilution, BioLegend) in PBS-BSA for 20 minutes at 4°C. Cells were washed twice with PBS-BSA and analyzed on a BD Accuri C6 flow cytometer. Binding signal was calculated by subtracting background fluorescence of cells incubated without sEFNB2-Fc and normalized across replicates based on total measured fluorescence for each day of experiments.

### Fluorescent Reduction Neutralization Test (FRNT)

Replication competent recombinant CedV (rCedV) chimeras were generated to display the G and F proteins of NiV_B_ and HeV and to encode a GFP reporter, as described previously (21, 37, 43). Vero 76 cells were seeded at a density of 2 × 10^4^ cells/well in black walled clear bottom 96-well plates (Corning Life Sciences). sEFNB2-Fc proteins were serially diluted 3-fold for 7-point dose response curves. An equal volume of Dulbecco’s modified Eagle’s medium with 10% v/v cosmic calf serum (DMEM-10) containing rCedV-NiV_B_-GFP, rCedV-HeV-GFP, and rCedV-GFP was added to each dilution for a final multiplicity of infection (MOI) of 0.05 and incubated for 2 h at 37°C with 5% CO_2_. Each of the virus-sEFNB2-Fc mixtures (90 μL/well) was added to the pre-seeded Vero 76 cells in triplicate and incubated for an additional 24 h at 37°C with 5% CO_2_. The supernatants of the virus-sEFNB2-Fc mixtures were removed, and the plates containing Vero 76 cells were fixed with 4% formaldehyde in PBS for 20 minutes at room temperature. The plates were then gently washed with diH_2_O and imaged using a CTL S6 analyzer (Cellular Technology Limited). Fluorescent foci were counted using the CTL Basic Count feature and S6 software. The 50% inhibitory concentration (IC_50_) was determined as the sEFNB2-Fc concentration at which there was a 50% reduction in fluorescent foci versus untreated control wells. The IC_50_ values were calculated by non-linear regression curved fit with variable slope with GraphPad Prism 9. The limit of detection for this assay was 50 fluorescent foci. Percent neutralization was calculated by normalizing counts to a virus only control.

## REAGENT AND DATA AVAILABILITY

Plasmids are deposited with Addgene under accession numbers 200975-200986. Illumina sequencing data are deposited in NCBI’s Gene Expression Omnibus (GEO).

## SUPPLEMENTAL MATERIAL

Supplementary Figures S1 to S7

Processed data files for deep mutagenesis of EFNB2-RBD in the wild-type and D62Q backgrounds

## ACKNOWLEDGEMENTS

Staff at the UIUC Roy J. Carver Biotechnology Center assisted with FACS and Illumina sequencing. Christine A. Devlin and Austin T. Weigle assisted with computational tools for preparation of heat maps. The views expressed in the manuscript are solely those of the authors, and they do not represent official views or opinions of the Department of Defense, the U.S. Government, the Uniformed Services University, nor the Henry M. Jackson Foundation for the Advancement of Military Medicine, Inc. Several of the authors are U.S. Government employees. This work was prepared as part of their official duties. Title 17 U.S.C. § 105 provides that ‘Copyright protection under this title is not available for any work of the United States Government.’ Title 17 U.S.C. §101 defines a U.S. Government work as a work prepared by a military service member or employee of the U.S. Government as part of that person’s official duties.

## FUNDING

D.S. acknowledges support from the National Institute of Health (Award No. R35GM142745). The development of chimeric henipavirus neutalization assays was supported by National Institutes of Health grant U19AI142764 to C.C.B.

## CONFLICTS OF INTEREST

E.P. is a shareholder and employee of Cyrus Biotechnology, which licenses and commercializes intellectual property held by the University of Illinois for soluble decoy receptors targeting SARS-CoV-2 and HCMV. Cyrus Biotechnology had no role in the design, execution, analysis or interpretation of the research described in this study.

